# Analysis of task-related MEG functional brain networks using dynamic mode decomposition

**DOI:** 10.1101/2022.06.08.495279

**Authors:** Hmayag Partamian, Judie Tabbal, Mahmoud Hassan, Fadi Karameh

## Abstract

**Objective:** Functional connectivity networks explain the different brain states during diverse motor, cognitive, and sensory functions. Extracting spatial network configurations and their temporal evolution is crucial for understanding the brain function during diverse behavioral tasks.

**Approach:** In this study, we introduce the use of dynamic mode decomposition (DMD) to extract the dynamics of brain networks. We compared DMD with principal component analysis (PCA) using real magnetoencephalography (MEG) data during motor and memory tasks.

**Main Results:** The framework generates dominant spatial brain networks and their time dynamics during simple tasks, such as button press and left-hand movement, as well as more complex tasks, such as picture naming and memory tasks. Our findings show that the DMD-based approach provides a better temporal resolution than the PCA-based approach.

**Significance:** We believe that DMD has a very high potential for deciphering the spatiotemporal dynamics of electrophysiological brain network states during tasks.

## 1. Introduction

Understanding task-related brain interactions have been studied extensively in recent years to quantify the changes in the electrical and magnetic activity of the brain and identify the active brain parts during specific tasks. The dynamics of these interactions evolve rapidly and vary, to some extent, between trials and between subjects [1]. However, knowing that the brain activity during a specific task is focal (localized at specific regions in the brain), specific brain regions are activated and communicate with other parts of the brain during visual [2], cognitive [3], motor tasks [4] among others. Most of these studies measure the magnetic and electric activity in the brain using non-invasive, high temporal resolution magnetoencephalography (MEG) and electroencephalography (EEG) techniques [5]. The time-series data acquired from MEG (or EEG) is processed to find sources within the brain from which the aggregated measurements are collected using beamforming methods [6] that are further processed to quantify the statistical interdependencies between different brain regions [7]. Different connectivity measures compute brain interactions using different assumptions such as the phase-locking value (PLV) that computes statistical phase correlations across trials [8] or the amplitude envelope correlation (AEC) [9] that computes amplitude correlations. To quantify the evolution of these networks across time, many model-free dimensionality reduction methods or clustering methods were used to extract data-driven networks. Temporal independent component analysis (ICA) was used to identify task-related functional networks from MEG data to track the dynamics of the brain networks during finger movement and memory tasks [10]. Principal component analysis (PCA) was also used to identify hidden coherent functional connectivity (FC) patterns in multiple sclerosis patients [11]. Other studies employed the non-negative matrix factorization [12] [13] and k-means [14] to learn dominant brain networks from EEG and MEG data. A recent study compared different data decomposition techniques to decipher the dynamics during three tasks: self-button press, fast finger movement, and working memory task [15], which showed globally modest performance of ICA/PCA-based methods.

On the other hand, dynamic mode decomposition (DMD) [16], is a modern data-driven powerful methodology to analyse nonlinear dynamics of time-series data which is a combination of PCA in space with Fourier analysis in time [17]. Originally designed to learn dominant spatial and temporal modes of activity in mechanical fluid dynamics [18], DMD has been used in many applications such as predictive maintenance of industrial time series data [19], infectious disease dynamical analysis [20], and neuroscience, mainly using fMRI [17] [21]. Specifically, coherent spatial and temporal patterns were identified in motor activation tasks and sleep spindles using DMD [17].

Cognitive brain responses exhibit transient changes that may vary from one trial to the other and between subjects. Across trials, averaging is a popular method used to increase the signal-to-noise ratio assuming that the event-related potentials are synchronous across the trials. However, inter-trial measurements are not stationary and some variability exists such as polarity shifts and temporal latency of the peaks that vary between trials and subjects. Therefore, inter-trail averaging may cancel out potentials that are not time-locked to the stimulus onsets or are shifted in polarity, and the resulting average will be distorted [1].

In this study, we employ the principal component analysis (PCA) and the dynamic mode decomposition (DMD) method to emphasize temporal similarities and approximate the dynamics of the networks by extracting data-driven spatial principal components (PC) with the corresponding temporal fingerprints and spatial and temporal DMD modes using MEG data recorded during 2 motor tasks and a memory task. We show that DMD which is robust to inter-trial and inter-subject temporal jitter variabilities can capture the dominant spatial and temporal brain configurations. The DMD results are compared with the PCA. Since the DMD modes are complex in nature, we evaluate the real, complex, and magnitudes of the modes to quantify the dynamical interaction between the different brain regions. In addition, to analyse the dynamics of longer recordings where multiple stimulations are applied, a sliding window approach is applied to extract local temporal principal components and DMD modes across trials and subjects that are further processed to find the dominant spatial configurations and identify the evolution of network states across time.

## 2. Materials and Methods

### 2.1 Data

#### 2.1.1 Dataset 1: Self-paced button press task

This dataset includes MEG recordings of 15 right-handed participants (9 male, 6 female, aged between 21 and 29). The participants were asked to use the index finger of their dominant hand once every 30 seconds. 275-channel CTF MEG system was used at a 600Hz sampling rate to acquire the data that were co-registered using subject-specific MRI which was parcellated using the AAL atlas with 78 regions [10] [24].

#### 2.1.2 Dataset 2: HCP left hand movement task

This dataset is extracted from the MEG data of 61 healthy individuals (28 male, 33 female, aged between 22 and 35) provided by the Human Connectome Project (HCP, MEG-1 release). Although the original data included the MEG recordings of left and right hand and foot recordings, only the left-hand movement trials were used in this study for simplicity. The MEG recordings were sampled at 508.6275 Hz and co-registered using subject-specific MRI [25].

#### 2.1.3 Dataset 3: Sternberg working memory task

This dataset includes MEG recordings of 19 healthy individuals (10 male, 9 female, aged 25±3years). The Sternberg task comprises two visual stimuli (geometric shapes showing on the screen for 0.6 seconds each separated by 1 second) followed by 7 second maintenance period. Next, another visual stimulus is shown on the screen and the subjects were asked to use their right index finger to press the button if the current image matches with one of the previously visual stimuli. Following this, an immediate feedback was given to show the correct response. For each individual, a total of 30 trials were recorded separated by 30 second rest periods using 275-channel CTF MEG system at 600Hz sampling rate and co-registered using subject-specific MRI [10]. The datasets were approved by the Research Ethics Committee of the University of Nottingham Medical School [10].

### 2.2 Methodology

#### 2.2.1 Processing Pipeline

In this section, we illustrate the processing pipeline adopted to analyse the brain activity during cognitive tasks and extract task-related dominant brain networks. Figure 1 describes the overall workflow adopted to build dynamic FC matrices as a preliminary step. First, the MEG data was collected during a specific task along with the subject-specific MRI data. After pre-processing the data to remove the unwanted noise and the artifacts, the MEG and the MRI data are used to estimate brain source activity using beamforming methods that project the external recorded MEG data inside the brain and estimate the time series (source signals) of *N*_*ROI*_ regions of interest (ROI). Next, the source data is processed to compute dynamic FC measures. In this study, we employ the AEC for the real datasets whose results are reported in the main document, and the PLV for the simulated data in the supplementary materials. Using *T* sliding windows, we generate a 3D tuple of size *N*_*ROI*_*xN*_*ROI*_*xT*. Due to symmetry, the upper triangular part of each connectivity matrix is extracted forming the 2D matrix 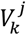 of size 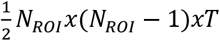 matrix for each trial *k* of each subject *j* that reduces the computational cost and redundancy in the data. Once all the matrices 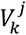 are computed for all trials per subject *j*, they are concatenated vertically to form the matrix 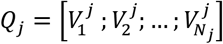 where *N*_*j*_ represents the number of trials for subject *j* as shown in figure 2.

**Figure 1.**
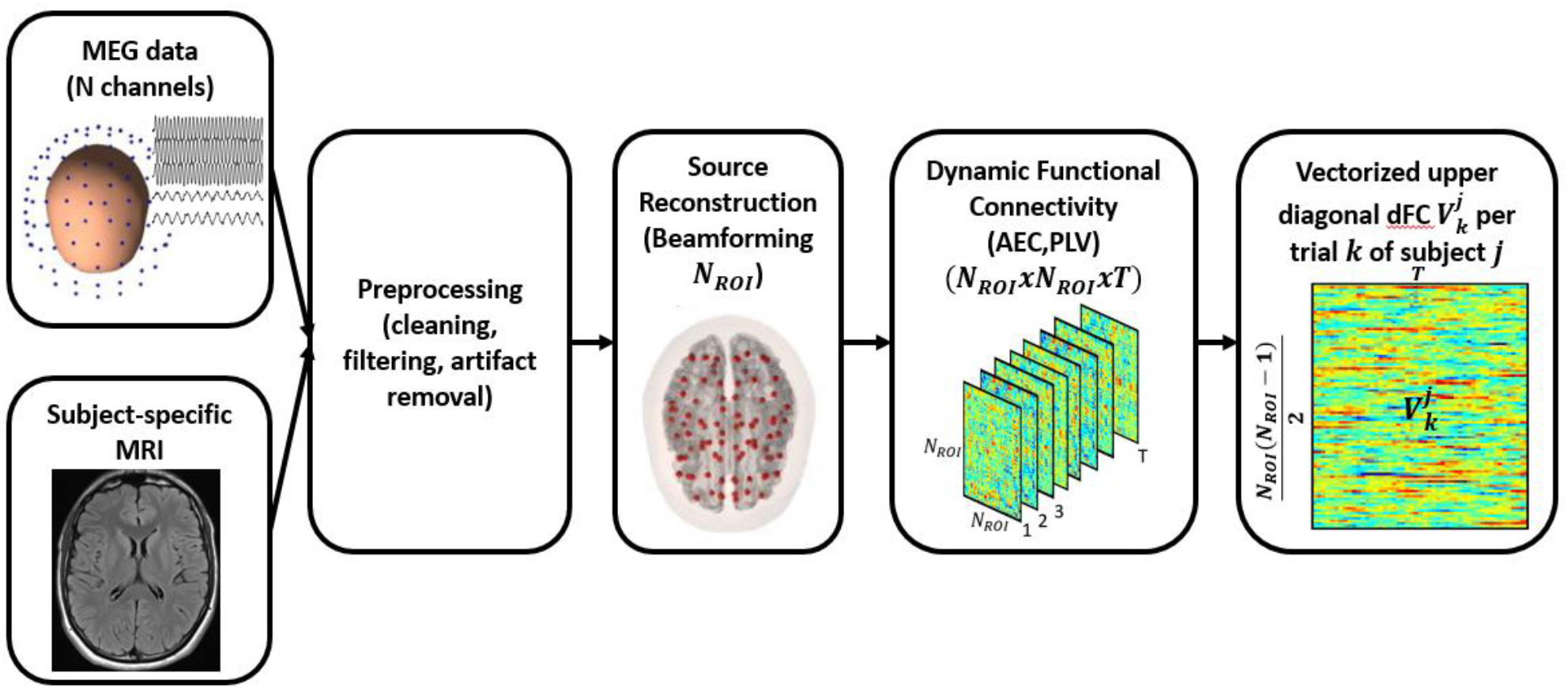
MEG data processing and computation of the FC during tasks per trial. The MEG data collected during a specific task along with the subject-specific MRI data is first processed to remove unwanted noise and artifacts. Next, source reconstruction is performed using beamforming method with *N*_*ROI*_ sources. Then, the source data is dissected into *T* windows and the amplitude envelope correlation connectivity is computed per window producing *T* dynamic FC matrices. These matrices are then processed to extract the upper diagonal part producing a vector for each window and are all collected into a single matrix 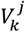.

**Figure 2.**
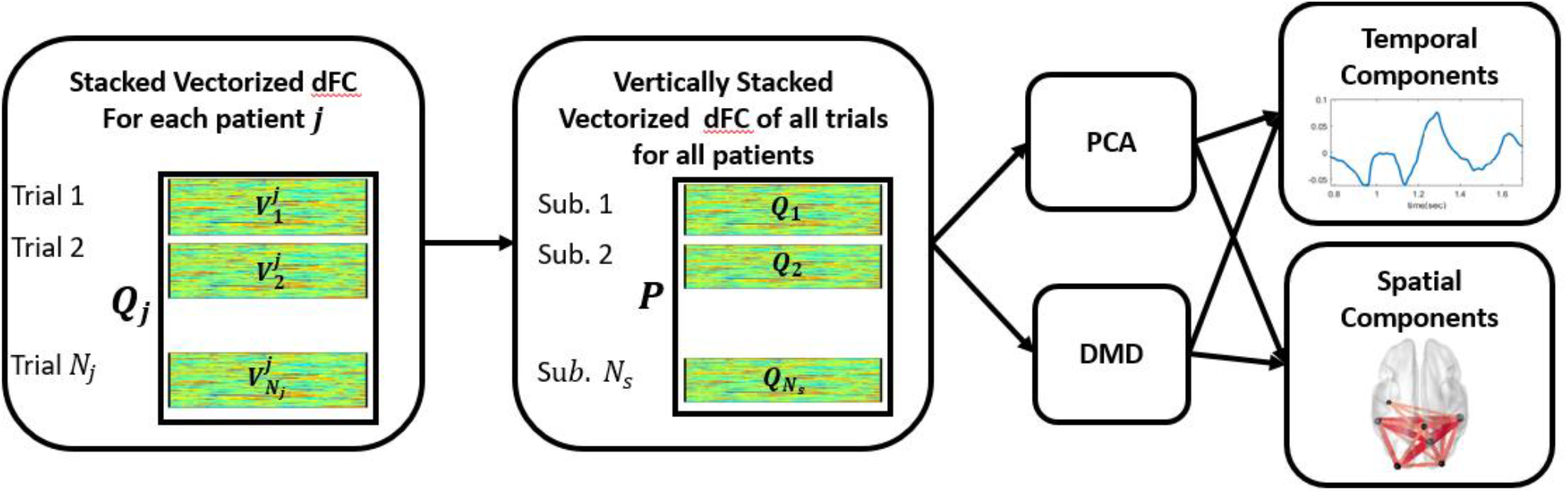
Dynamic analysis of a window of FC during a task across subjects using signal decomposition techniques. The vectorized dynamic FC matrices of all the trials are stacked vertically for each subject ***j*** into a matrix ***Q***_***j***_. Similarly, all the matrices ***Q*** for all subjects are concatenated vertically to form a matrix ***P***. Next, PCA and DMD are applied to learn dominant temporal and spatial components.

Next, the matrices *Q*_*j*_ are concatenated vertically for all subjects *N*_*s*_, to form the final matrix *P* which is then processed using PCA and DMD methods to find temporal and spatial principal components and modes. The extracted vectorized spatial components which represent the upper triangular part of the connectivity matrices are made symmetric to form the complete connectivity matrices. The spatial components of PCA and the modes of DMD represent the spatial brain networks while the temporal components and the temporal modes represent the time dynamics of the network evolution. In addition, the spatial networks are statistically analysed using t-test with >95% confidence value with Bonferroni corrections compared to the null distribution [26]. While simple tasks encompass a few network configurations, more complex tasks exhibit dynamic evolution of networks in the brain which need to be analysed with higher temporal resolution. As shown in figure 3 A, instead of processing the whole dFC data *P* for the whole interval, the data is dissected into shorter windows of size *w* and the signal decomposition methods (PCA, DMD) are applied dynamically using overlapping sliding windows on the dFC matrix. Since the complexity of the task is usually unknown, this approach is a generalized form of the method that allows the study of dominant networks when the dynamics of the network change rapidly across time. In this study, this approach is used for the working memory task only (as well as the simulated data in the supplementary material). For each window, the stacked vectorized data is processed and a small number of components and modes (e.g. 2 or 4) is extracted assuming that a few network configurations are only present within a specific time interval. After all the windows are analysed and dominant networks are extracted, they are all stacked and PCA is applied to find the dominant networks and their respective temporal evolution.

**Figure 3.**
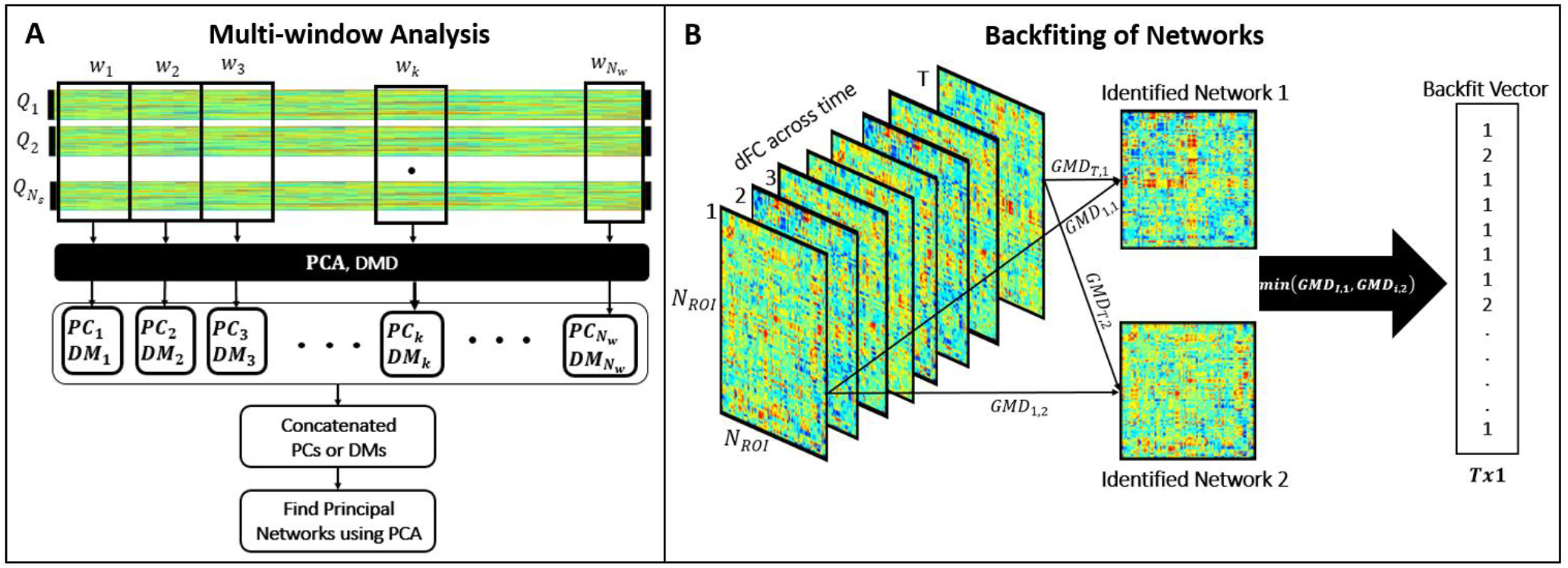
A. Multi-window dynamic analysis of FC across subjects for long-duration tasks. In general, when analysing tasks with multiple stimulations, the procedure is upgraded to study windows of FC time series. First, we dissect the data dFC data into windows and learn the dominant spatial configurations (PCs and DMs) following the procedure described in figures 1 and 2. Then, all the extracted dominant networks are concatenated and PCA is applied to find the dominant networks across all windows along with the temporal components to identify timestamps for each network. **B. Backfiting Algorithm**. dFC matrices across time are compared with the identified networks based on global map dissimilarity measure. The label of the minimum GMD with the two candidate networks is inherited by the dFC resulting in a backfit vector of size ***T***.

### 2.3 Data Pre-processing

Datasets 1 and 3 were pre-processed and bad segments from muscle, eye, and head movement were visually examined and removed as described in [10]. Dataset 2 was pre-processed according to the steps described by the HCP consortium. Bad channels, noisy segments, and unwanted independent components were removed. Then, each dataset was segmented and filtered according to the performed tasks.

Segments were extracted in the interval of [-12; 12sec] relative to the button press onset for dataset 1, and in the interval of [-1.2; 1.2sec] relative to the electromyogram (EMG) onset for dataset 2. MEG data of both datasets 1 and 2 were filtered in the beta band [13-30Hz] to keep the frequency components of interest relative to motor tasks. Regarding dataset 3, we segmented trials in the interval [-16; 28sec] relative to the stimulus presentation and filtered data in the [4-30 Hz] bandwidth to keep all the frequency components relative to the task.

The MEG data collected in datasets 1, 2, and 3 were processed to localize brain sources and estimate their activities using linearly constrained minimum variance beamforming (LCMV) on parcellated cortex using the AAL atlas with 78 ROI and by employing the subject-specific anatomical MRI registered to an MNI template. This procedure was followed by an inverse registration to project the data onto the anatomical subject space the details of which can be found in [15]. Note that all trial data was used for the self-paced button press task with a total of 510 recordings for all the participants combined. Since the number of subjects for the HCP data was too large (61 subjects) each with a very large number of trials (total of 9150 trials), averaging across trials per subject was used to avoid memory issues associated with the PCA computation. Therefore, we use the 61 averaged dFC data for the evaluation of the HCP data.

### 2.4 Functional Connectivity Measures

Next, the dynamic functional connectivity (dFC) matrices were computed. In this work, we employed two FC measures as follows. AEC using the Hilbert transform between all ROIs was applied for datasets 1 and 3 followed by leakage correction (multivariate symmetric orthogonalization [27]) to avoid erroneous estimates of FC matrices. To account for the dynamic behavior of FC estimates, we used the sliding window approach for both datasets 1 and 3 where 6-second intervals with 0.5-second overlap were adopted. Since the duration of the recordings of dataset 2 was relatively short, the instantaneous amplitude correlation (IAC) measure that extracts high temporal resolution amplitude envelope correlation, was computed for each trial of dataset 2 followed by pair-wise orthogonalization since the interval is relatively short [28][29][30] for leakage correction. Finally, PLV was computed across trials per subject for the simulated dataset whose results can be found in the supplementary material.

### 2.5 Source Separation techniques

In this study, we employ two signal decomposition techniques. PCA is a statistical procedure that computes orthogonal principal components that explain most of the variance [31][32]. The components are ranked from the one which explains the highest variance in the data to the least. Our aim here is to employ the FC time series of the source data to find principal spatial components by emphasizing temporal correlations. Thus, we employed vertical concatenation of the data across trials and in all subjects to find dominant spatial configurations that behave similarly across time.

On the other hand, DMD is another dimensionality reduction method that computes the coherent structures from the raw data in the form of modes that contain spatial information associated with an oscillatory temporal evolution at a fixed frequency that could be pure, decaying, or growing. The modes and the eigenvalues computed are approximations of the Koopman operator [33]. Although DMD relies on PCA to extract the coherent structures in the data, however, DMD is different from PCA since the modes are not necessarily orthogonal and the temporal behavior is often predetermined to be oscillatory which can be more interpretable. This is true because DMD finds the best linear operator in the mode space that fits the data and links it back to the channel or node data through the DMD mode matrix.

A detailed description of the PCA and the DMD approaches is available in the supplementary material.

### 2.6 Choice of Parameters

The identification of the optimal number of components for each task is challenging. Usually, this value is determined according to a certain percentage of variance explained in the data in the case of the PCA and percentage reconstruction in the case of the DMD, however, in this study our aim is to find only the significant components. The DIFFIT method [15] which computes the differences in data fitting while varying the number of components *r*, can help identify the number of components in the case of the PCA and the number of modes in the case of the DMD, that produce the highest fit difference. The value of *r* was varied from 2 to 10 and the value with maximum DIFFIT magnitude was chosen. On the other hand, the *r* networks identified are further processed using the *t*-test hypotheses testing to identify the regions that are outliers to the assumed Gaussian distribution. Here, *r* represents the number of components for the PCA and the number of modes for the DMD. DMD can be further smoothed by Hankel time delay embeddings which enriches the data by concatenating vertically *h* time delayed snapshots of the data as detailed in the supplementary material. We employ *h* =2 time delay embeddings since the condition to capture the full dynamics is already met due to the large number of trials per study.

### 2.7 Backfiting

The objective of this step is to analyse the obtained group-level states at trial and subject levels. The identified dominant networks are used to fit the microstate templates back to the original data networks. Therefore, the dFC data across time is compared and matched with one of the identified principal networks [34]. The method calculates the global map dissimilarity (GMD) measure [35] of each time-stamp with the identified components and labels each dFC to its matching microstate. As an example, consider figure 3B where the dFC matrices are labelled temporally as 1,2, … *T*. The GMD of each dFC is computed with each of the identified networks 1 and 2, and then the minimum is used to label the dFC network as 1 or 2. Therefore, the resulting backfit vector contains a label for each of the *T* dFC networks.

### 2.8 Statistical Significance

The average statistic is not an adequate measure to quantify multiple trials and subjects. Therefore, we employ the spatial components generated by the PCA and the DMD algorithms for all the trials and apply the *t*-test with 95% confidence interval with Bonferroni corrections to find statistically significant network components. Having 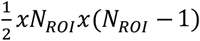 connectivity vectors for *T* timestamps for each trial, PCA outputs a vector of size 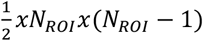 for each principal component, each representing connectivity between two channels. Each of the connectivity values is statistically evaluated separately across all trials and all subjects. First we compute the mean and the standard deviation of all the data and assume our null hypothesis to be a Gaussian with the computed mean and variance. Next, each component of the dFC vector is considered to be a sample from the population. The t-test is applied to identify the connectivity components that are rejected at 95% confidence interval. Bonferroni corrections are applied since we are performing multiple comparisons across all the source nodes.

## 3. Results

To provide a comparison between PCA and DMD, we provide in the supplementary material a comparative analysis on synthetic toy data to pinpoint the effect of inter-trial and inter-subject variability in terms of temporal phase lags. The assumption is that different trials (subjects) exhibit different responses during the recording process. Thus, we present the analysis of the toy data example to study how DMD and PCA respond to temporal latency jitters in the supplementary material. Briefly, the results show that DMD is robust to the temporal latency jitters and can capture the spatial and the temporal dynamics more robustly when compared to the PCA. In addition, the supplementary material contains results on simulated EEG data [22] as a proof of concept.

In this section, we report the results of the methodology described in section 2 on three real MEG datasets to extract dominant spatial brain configurations and their respective temporal dynamics.

To determine the number of components to consider, we employ the DIFFIT technique. For the self-paced button press and the left-hand movement datasets, the DIFFIT function found that 2 components were enough to describe the spatial configurations. Finally, for the working memory task dataset, we computed the DIFFIT values across all the windows and in each window, the optimal number of components was found to be 2 as well. Note that, since we are localized in time in our computations and we are stacking the trials and subject data preserving the temporal localization, the results of the DIFFIT function (which is 2 in all cases) is valid since brain networks can ideally have a few configurations at a time. Thus, we employ 2 components in all our studies. In the case of the DMD-based analysis, since 2 modes correspond to one, we employ 2 modes only but use the real and the imaginary networks as separate networks. We will be discussing this phenomenon in the experiments below. For the first 2 real data sets, the results show the mean spatial networks, the temporal dynamics, the statistical networks, and the spatial network similarities across all subjects and their trials across time using backfiting. Since the first two datasets were processed to extract dFC for 49 and 36 time stamps respectively, we employ the workflows depicted in figures 1 and 2. However, for the case of the memory task we employ additionally process the data with the workflow stage depicted in figure 3 where stages 1 and 2 of the pipeline are applied in a sliding window fashion and then a global PCA is applied to find the common spatial configurations and their corresponding temporal evolution time series. The motor task results show the dominant networks and the corresponding time stamps with color-coded boxes that indicate dominant structures at specific intervals that are compared based on the stimulations applied during data acquisition.

Note that for illustration purposes, we manually tuned the thresholds for the mean networks (figures A and B in figures 4 and 6 and figures A, B, and C for figures 5 and 7) in the following manner: 75 % of the maximum for the self-paced button press task and the HCP task, and 65 % for the working memory task. To avoid the thresholding variability, we added the networks after statistical analysis with *t*-test at 95% confidence interval with Bonferroni corrections (in E and F of figures 4 and 6 and F,G, and H of figures 5 and 7). Finally, the results show the backfiting results across all trials for the first two datasets and the global PCA temporal components for the working memory task.

**Figure 4.**
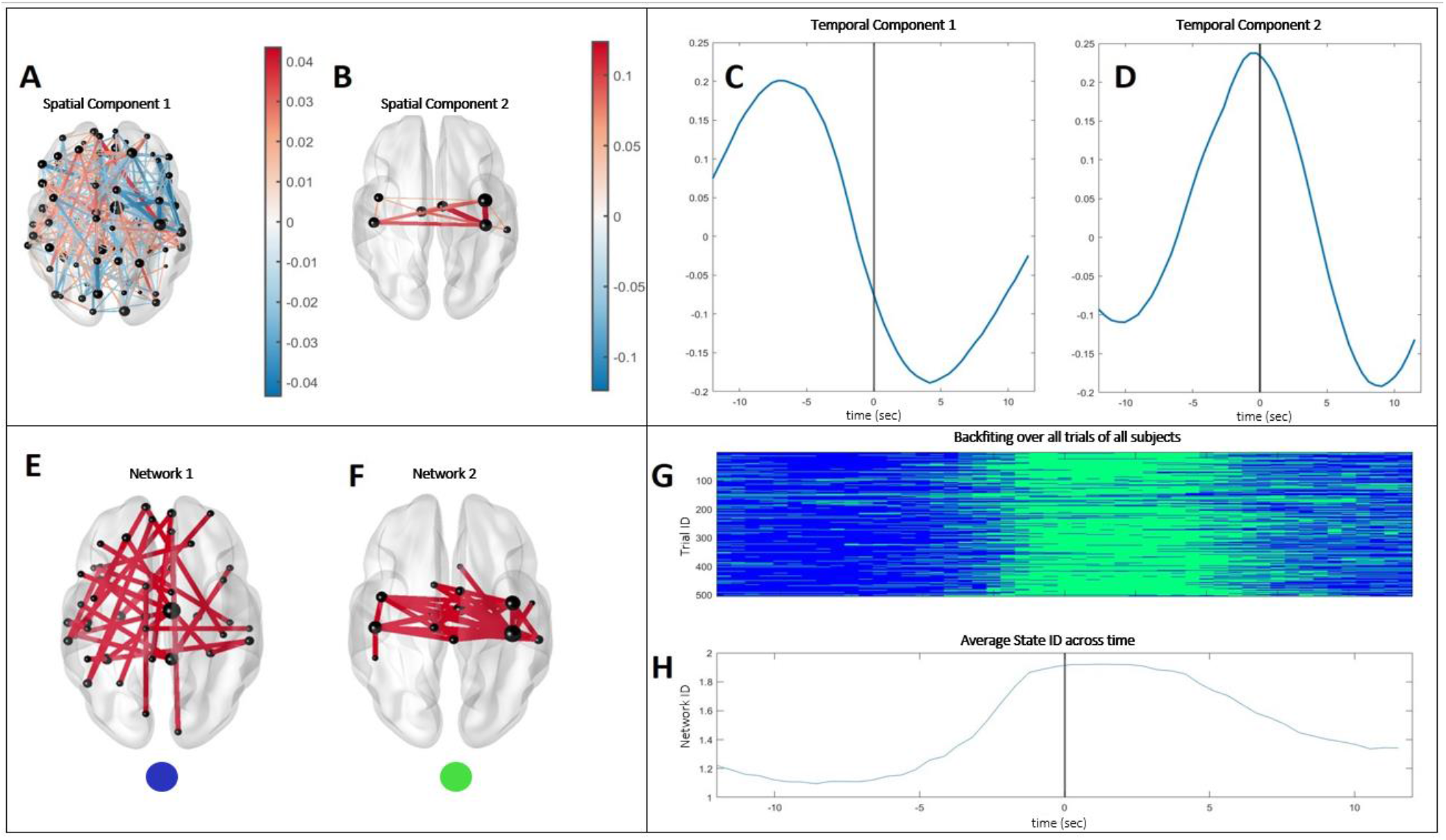
Self-paced button press task networks using PCA. A and B represent the dominant spatial components identified using the PCA approach. While network A is random, network B shows a motor network in the central region of the brain. C and D represent the corresponding temporal components which shows where the network is dominant across time. Figures E and F represent the networks identified using 95% confidence interval using the ***t***-test with Bonferroni correction. G depicts the results of the backfiting algorithm when spatial components 1(blue) and 2 (green) are used as dictionary elements. H represents the averaged network ID across time which shows that the motor network 2 (green) is dominant during the button press when averaged across all trials of all subjects.

**Figure 5.**
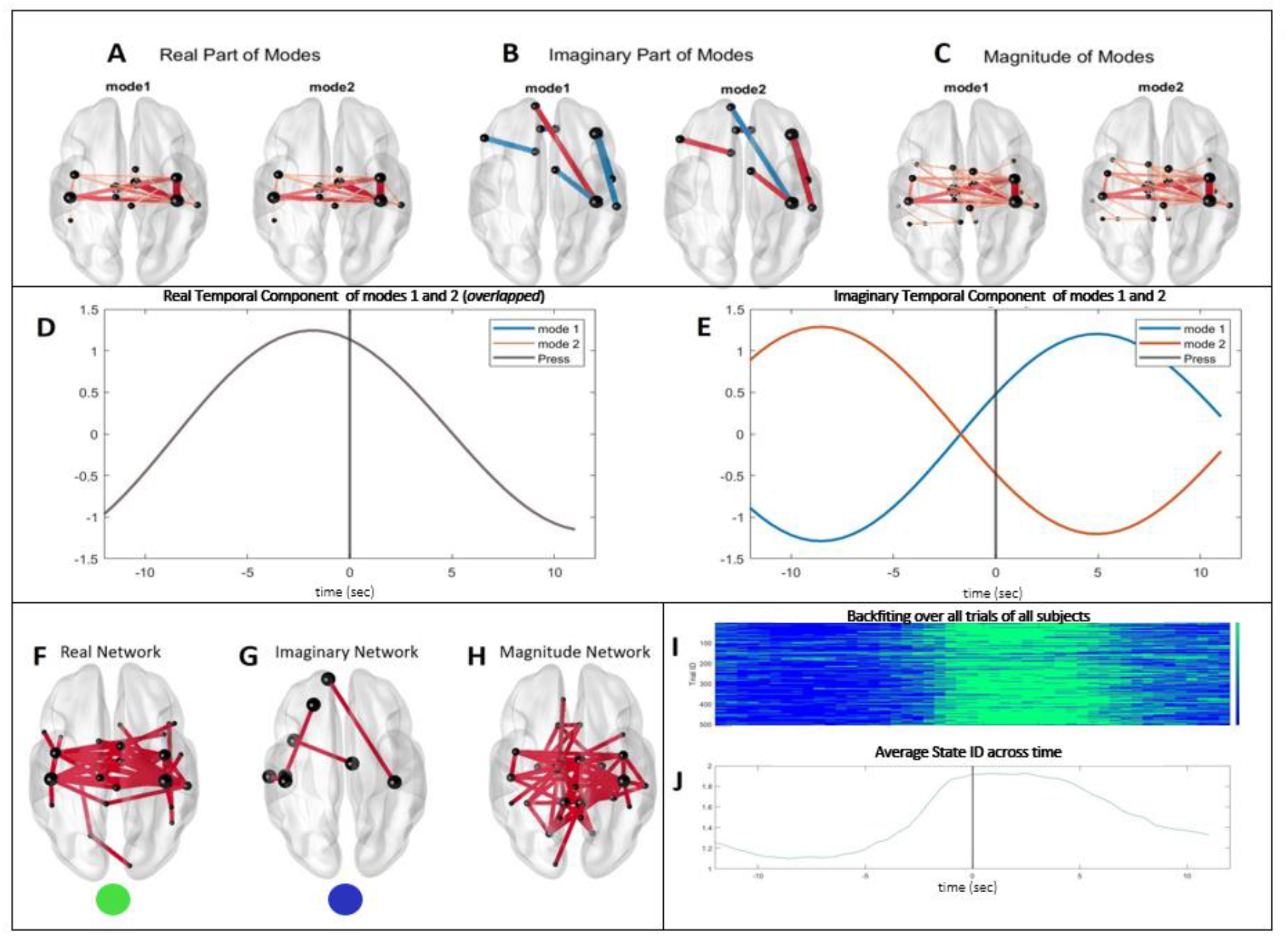
Self-paced button press task networks using DMD. A represents the real part of the identified modes which shows the motor task identified earlier using the PCA, B represents the imaginary part and C represents the magnitude of the identified networks which is equivalent to the motor network. D and E represent the corresponding temporal components which show where the network is dominant across time. Note that the real part of the time dynamics of the 2 modes are overlapping (D) and the imaginary parts are at **90°** phase difference. Figures F, G, and H represent the networks identified using 95% confidence interval using the ***t***-test with Bonferroni correction. Figure I depicts the results of the backfiting algorithm when the real spatial components (blue) and the imaginary spatial component (green) are used as dictionary elements. Finally, H represents the averaged network ID across time which shows that the real network which corresponds to the motor network (green) is dominant during the button press when averaged across all trials of all subjects.

**Figure 6.**
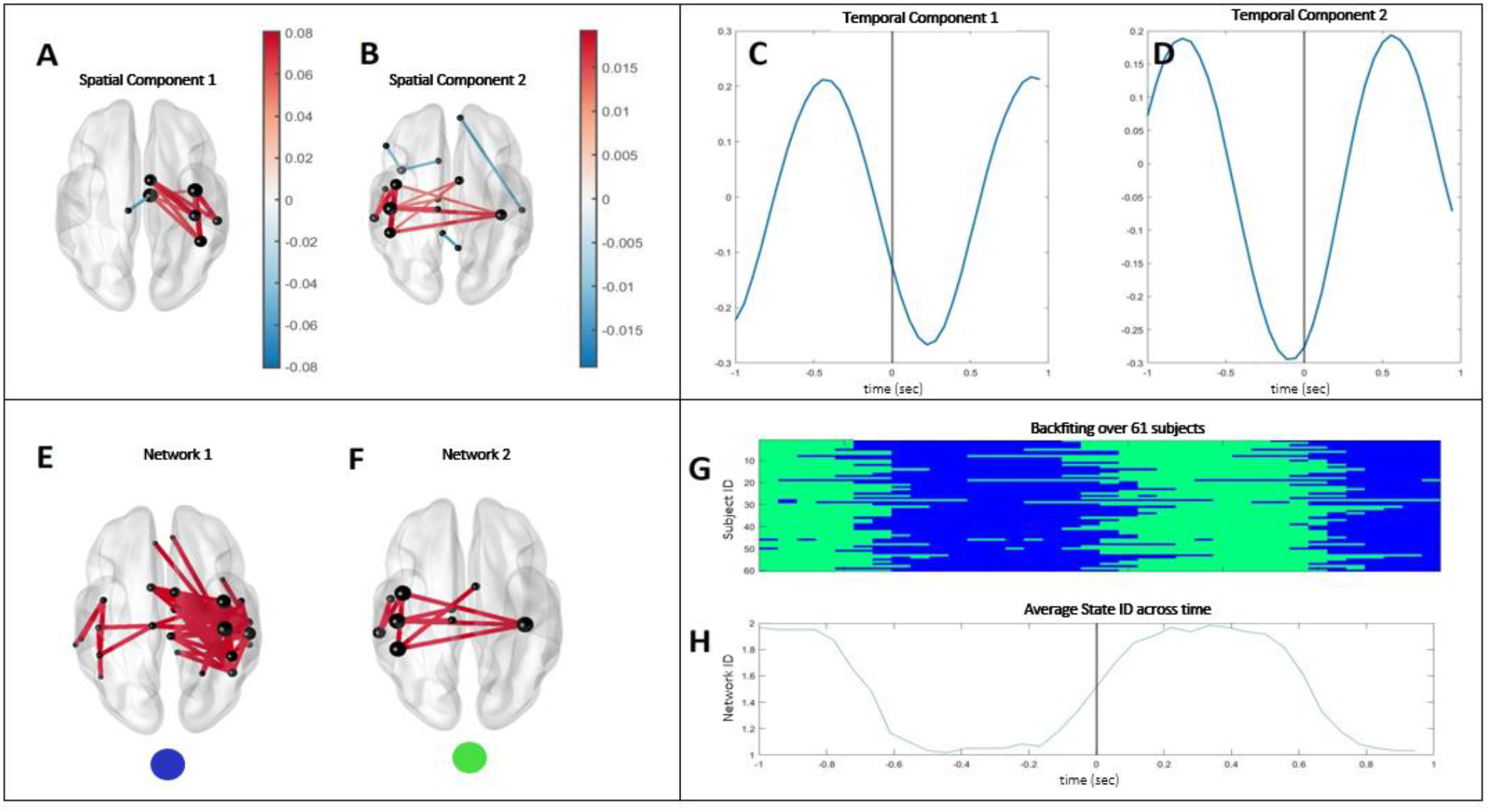
HCP networks using PCA. A and B represent the dominant spatial components identified using the PCA approach. A shows a right central network while B shows a left central network. C and D represent the corresponding temporal components which show where the network is dominant across time. Figures E and F represent the networks identified using 95% confidence interval using the ***t***-test with Bonferroni correction. G depicts the results of the backfiting algorithm when spatial components 1(blue) and 2 (green) are used as dictionary elements. H represents the averaged network ID across time which shows that network 1 (blue) is dominant prior to and during the button press when averaged across all trials of all subjects.

**Figure 7.**
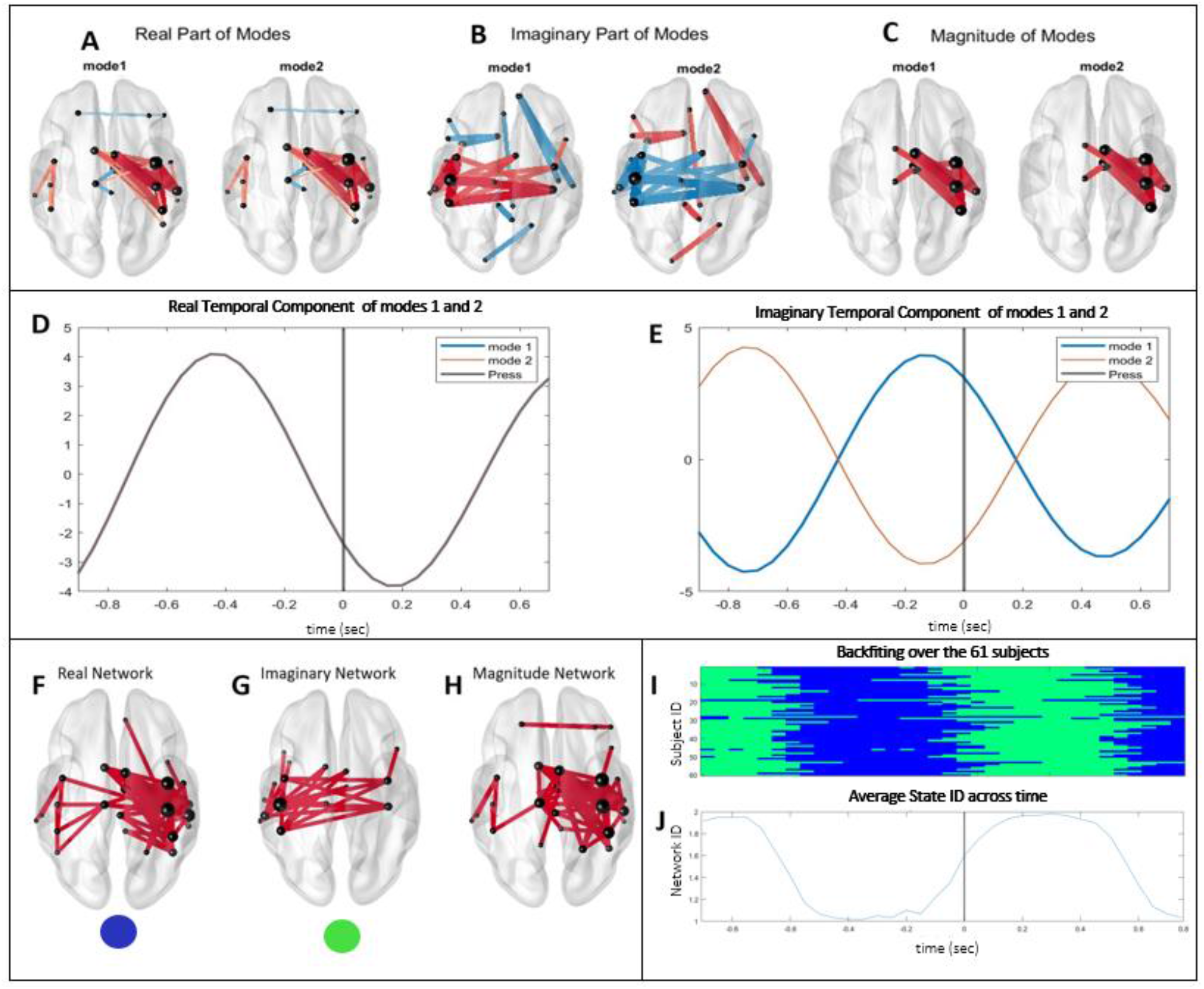
HCP networks using DMD. A represents the real part of the identified mode whose activity is predominantly focused in the right central region of the brain, B represents the imaginary part (left central activity) and C represents the magnitude of the identified network which only shows the right motor network identified in the real part. D and E represent the corresponding temporal components which show where the network is dominant across time. Note that the real part of the time dynamics of the 2 modes are overlapping (D) and the imaginary parts are at **90°** phase difference. Figures F, G, and H represent the networks identified using 95% confidence interval using the ***t***-test with Bonferroni correction. Figure I depicts the results of the backfiting algorithm when the real spatial components (blue) and the imaginary spatial component (green) are used as dictionary elements. Finally, H represents the averaged network ID across time which shows that the real network (blue) is dominant prior and during the button press when averaged across all trials of all subjects.

### 3.1 Self-Paced Button Press data

First, the self-paced button press data was analysed using the workflow shown in figures 1 and 2. The 24-second data was processed using a sliding window approach to extract 49 dFC matrices. The upper triangular parts of each dFC were extracted and processed using PCA to extract the two dominant networks. The number of components was determined using the DIFFIT method discussed in section 2. Note that, with the vertical concatenation, the PCA outputs common temporal dynamics across all the data and a set of spatial components for each trial. To understand the spatial distribution of the dynamics we employ two methods.

First, the components are averaged across all trials to find the two principal spatial networks. The first spatial network which is shown in figure 4A shows a random network that is dominant before and after the button press (at the 0-second mark). On the other hand, the principal network 2 shown in figure 4B shows activity restricted to the central region of the brain and is identified as a motor network since it occupies the motor cortex (that includes the precentral, paracentral, and the rolandic) and somatosensory cortex (which includes the postcentral, parietal and supramarginal areas). This network is dominant during the button press task as shown in figure 4D where the components are maximum around the 0-second mark at which the button was pressed. Figures 4E and 4F show the networks with the *t*-test at 95% confidence intervals. Employing the two networks as the elements of the dictionary microstates specific for this task, the backfiting algorithm is used to find the intervals of time during which each of them occupies across all trials of all the participants.

Note that, the dataset which comprises data from 15 subjects has a variable number of trials per subject which adds up to 506 trials. This allows us to compare the different trials’ and different participant’ brain configuration evolution during this simple button press task. As shown in the figure 4G, principal network 1 (the random microstate color-coded in blue) occupies the intervals of time before and after the button press in the majority of trials across all subjects while the principal network 2 (the motor microstate color-coded in green) occupies the interval during the button press in all the trials. Figure 4H is the averaged value of the state numbers across all trials of all subjects which clearly shows the dominance of the motor network during the button press task since its value is close to 2. Similar to the PCA-based analysis, DMD is applied to identify the dominant networks. In this case, DMD extracts complex-valued modes and time dynamics. Note that, although the time dynamics is unique for all the trials and subjects per mode, the spatial components embedded in the DMD mode matrix Φ are not i.e. each trial contains its corresponding block in the Φ matrix. Figures 5 A, B, and C show the average of all the components of all trials and subjects for the real, imaginary, and magnitude of the first two modes. The corresponding unique real and imaginary parts of the time dynamics are shown in figures 5D and E. Note that the two modes have overlapping real time dynamics and 90^0^ shifted imaginary temporal dynamics which is within the nature of the DMD decomposition as detailed in the supplementary material. Figures 5F, G, and H show the networks with the *t*-test at 95% confidence intervals.

The results are similar to the PCA-based method where the motor task dominates during the button press task. Considering the real and the imaginary networks as the principal microstates, back-fitting across all trials of all subjects also shows the inter-trial similarity across time where the motor network dominates the interval during which the button is pressed as shown in figures 5I and 5J. We can conclude that both PCA and DMD approaches successfully extracted a motor network during button press. DMD was also able to find a modulated network with the motor network in the left frontal region.

### 3.2 HCP left hand movement data

Similar to the self-based button press processing, the HCP left-hand movement data was analysed using the workflow shown in figures 1 and 2. The 2-second data was processed using a sliding window approach to extract 36 dFC matrices. The upper triangular parts of each dFC were extracted and processed using PCA to extract the two dominant networks which were averaged and statistically processed to identify them. Figure 6 shows the extracted dominant spatial and temporal components. The first mean network shown in figure 6A represents right central brain activity. Knowing that the task was moving the left hand, the dominant network identified here is justified since the right central cortex of the brain controls the left-hand movement. The connectivity drops just after the button press as seen in figure 6C. On the other hand, figure 6B shows another network that is active in the center left part of the brain, whose temporal dynamics indicate its presence before and after the button press as shown in figure 6D. The network components shown here were thresholded manually at 80%. We also analyse the principal components across all the subjects statistically using the *t*-test. The resulting networks for both principal components are shown in figures 6E and F. Next, back-fitting is applied across all the subjects using the two detected dominant components as microstates across all subjects (on the averaged dFC matrices across trials resulting in 61 samples) which shows the dominance of the first spatial component before the hand movement across all subjects as shown in 6G and H. Next, instead of using the PCA, we apply the DMD and identify the real, the imaginary, and the magnitudes of the first two dominant modes. As seen in figure 7, the real part of the DMD mode represents the right cortex motor activity and the imaginary part represents the left motor cortex network identified with the PCA analysis as a separate component. However, when the magnitude is computed, the right cortex network dominates and masks the left cortex movement. These results suggest that the identified networks are synchronous in time and are modulated together. The results also indicate that the left side activity of the brain is smaller in magnitude than the right side. This is true because DMD’s real and imaginary components are out of phase with one another. While the PCA identified the components separately, DMD identified them in a single mode. Figures 7D and E show the dynamics of these components in time that match the PCA results as well.

Finally, we employ the components of the DMD matrix Φ corresponding to all the subjects and apply the *t* −test to find statistically significant network components similar to the previous task. Considering the 61 components of the Φ matrix as different samples, the significant real, imaginary, and magnitude networks are shown in figures 7F,G, and H respectively. The real part and the imaginary part of mode 1 are taken as the microstate dictionary elements and backfiting is applied across the 61 subjects which shows similar results as the PCA as shown in figures 7I and J. The dominant right central motor network identified in this study shows that it is dominantly occupying the time interval before the button press. With the DMD-based approach, we can also see that the left central network (identified as the imaginary part of the DMD spatial mode) is modulated with the real network.

### 3.3 Working Memory Task

During complex tasks, different active parts of the brain change rapidly over time. Therefore, based on previous studies, it is expected to identify visual, semantic, sensorimotor, and other networks [15]. However, these networks shift rapidly and decisions to press a button if the third stimulus matches one may not be identified when processed in a batch across all trials and all subjects. The number of dFC networks computed is 75 windows in which the dynamics change rapidly. Therefore, we process the data in a sliding window approach to analyse using *w* =10 timestamps per window with 90% overlap. We extract dominant networks in each window to summarize the active networks across all trials and subjects and then identify dominant networks and their evolution across time using PCA based on the workflow depicted in figure 3. We extract two dominant components per window across all subjects for the PCA-based approach and 4 dominant modes (equivalent to 2) in the case of the DMD-based method where the magnitude networks are considered only. Next, PCA was applied to find 4 dominant networks across time.

Figure 8 shows the principal components of the PCA-based method. PC1 which represents the visual network (occupying mainly the occipital areas, cuneus, calcarine, and lingual brain regions) is the most dominant and recurring across time. The other identified networks are the semantic (bilateral parietal and temporal areas), the sensorimotor (the motor cortex), and an auxiliary network. Inspecting the temporal dynamics of each network, it can be observed that the visual network (blue line) is high before the 0-second mark (the first stimulus) and during the second stimulus as well as around the 5-second timeslot and finally during the probe stimulus (around the 10-second mark) and later around the 14-second mark. On the other hand, PC2 which represents the semantic network is only high (red box) during the semantic processing phase between the 2^*nd*^ stimulus S2 and the probe stimuli S3. The other networks are not identifiable.

**Figure 8.**
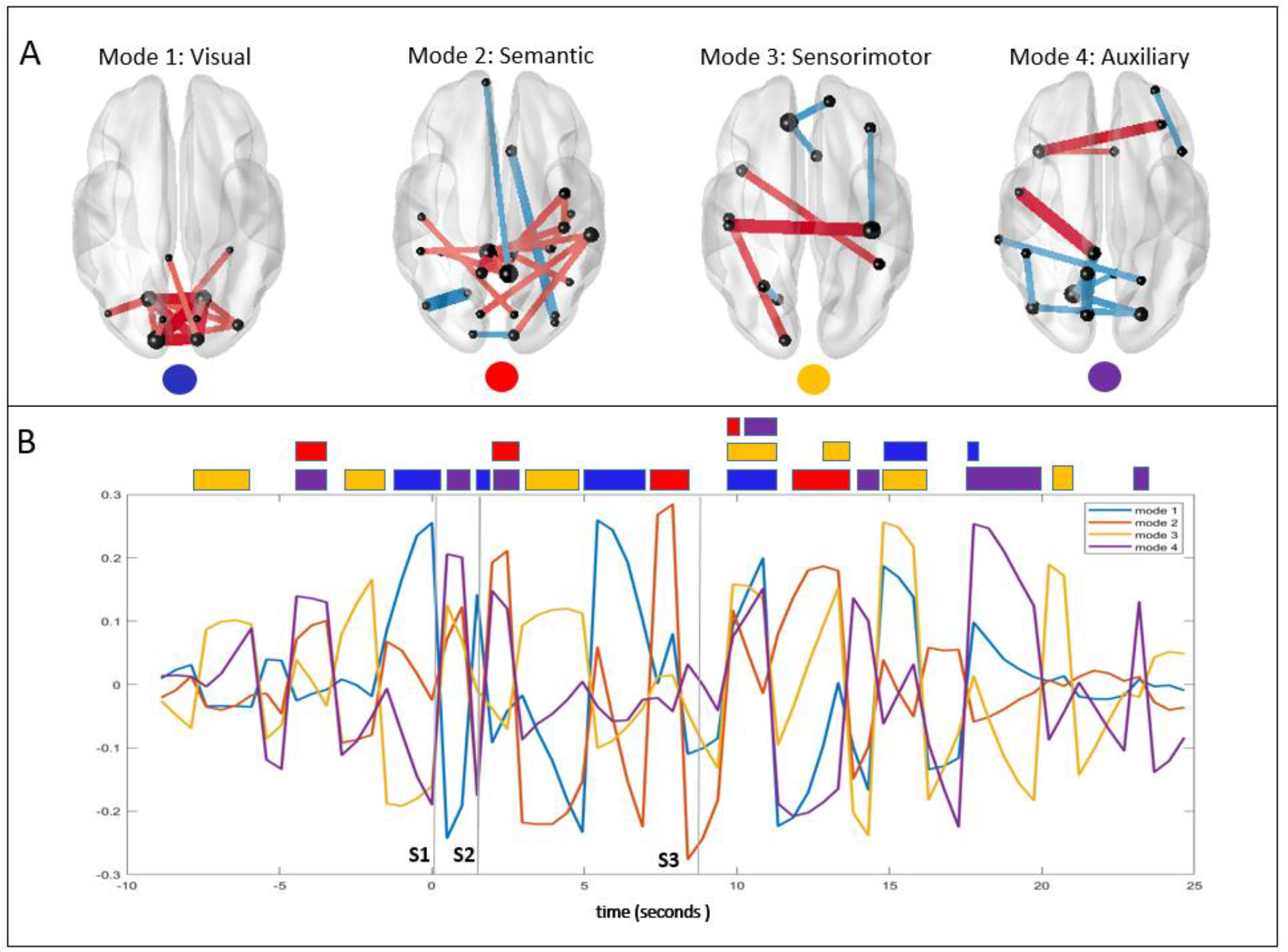
Working Memory Networks using PCA-based approach. A shows the four networks identified that were classified as visual (blue), semantic (red), sensorimotor (yellow), auxiliary (purple). B shows the temporal evolution of these networks with the corresponding color codes. At the top of the figure, the dominant network color codes show when the networks are mostly active.

Similarly, figure 9 shows the identified networks of the memory task for the DMD-based approach where the visual network is clearly expressed during all the visual stimuli as indicated by the blue boxes. On the other hand, the semantic network is active before the probe stimulus S3. In addition, the sensorimotor network can be seen to be active during the first stimulus and after the probe stimulus (yellow box). In addition, when compared with the PCA-based approach, the network configurations have a smoother and more localized in time than that of the PCA-based approach. We can see that DMD-based approach offered better temporal resolution when compared with the PCA based approach.

**Figure 9.**
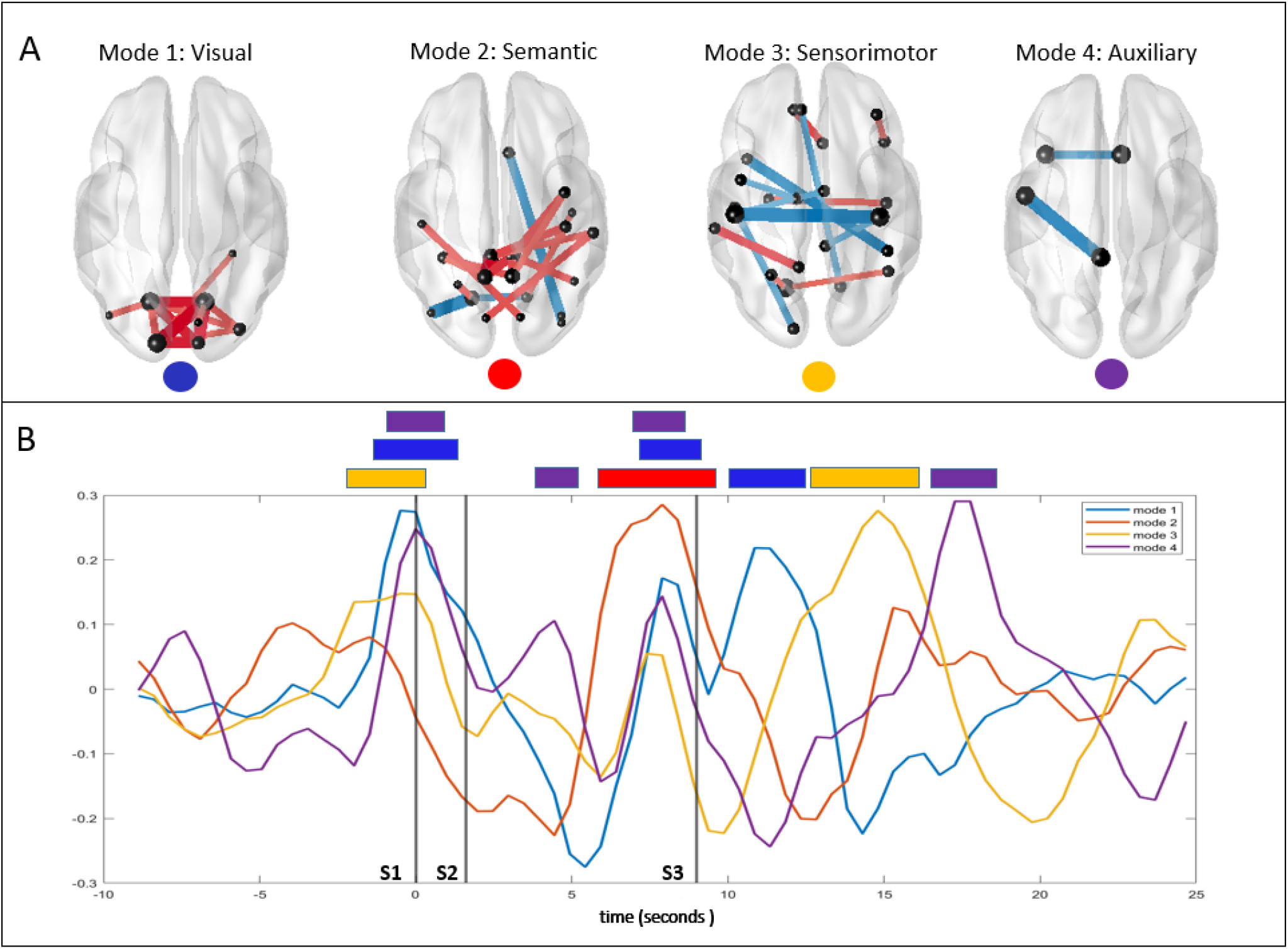
Working Memory Networks using DMD-based approach. A shows the four networks identified that were classified as visual (blue), semantic (red), sensorimotor (yellow), auxiliary (purple). B shows the temporal evolution of these networks with the corresponding color codes. At the top of the figure, the dominant network color codes show when the networks are mostly active. With the DMD-based approach, unlike the PCA-based approach, the networks occupy sparse intervals across time.

## 4. Discussion and Future Work

In this study, we introduced the DMD methodology and compared it with the PCA method to automatically extract spatial components and the temporal evolution in three different tasks. Our methodology employs all the trials in the data and imposes temporal constraints by processing the stacked trial data of all patients all at once taking inter-trial and inter-subject variability into consideration.

In summary, our work employs the PCA and the DMD method to extract dominant brain network configurations and the corresponding time dynamics. The resulting networks per trial are then averaged and statistically analysed to find significant spatial networks. Since the length of the datasets are different and brain networks often change states rapidly, we analysed dynamic networks using a sliding window approach. Overall, our results show that both PCA and DMD approaches successfully extracted dominant spatial configurations and their respective time dynamics on the simulated and the real MEG data when compared with the results of previous works that used the same dataset [15] [23]. Simulated data results that were reported in the supplementary material showed how network configurations shifted across time and were comparative with the ground truth. Although the two methods showed different network configurations due to the relative mathematical assumptions, the results were overall consistent. For the self-paced button press task, the dominant network identified with both PCA and DMD approaches was the motor network that was dominantly active during the button press across all trials and subjects. For the left-hand movement task, the dominant network was found to be the right central hemisphere, which is justified since the right hemisphere controls the left side limb movements. Another network was also identified on the left central region with smaller magnitude. With the DMD approach, both these networks were embedded into a single mode which suggests that the two networks synchronously turn on and off together. This is true because the DMD real and imaginary parts are often at 90 degrees phase shift. In the PCA-based results, one can see this phase shift in the time dynamics as well. Therefore, the left-hand movement activates the right and left motor networks in an alternating fashion. Finally, the complex motor task data was evaluated and the results show a dominant visual network active across time in all trials and subjects. The visual network dominated during the first stimulus and before and after the probe stimulus S3 similar results in terms of localization in time were reported in [15]. The methodology also identified a semantic network that had the highest activity before the probe stimulus S3. The sensorimotor network identified peaks before the first stimulus S1 and the probe stimulus S3 although in [15] the sensorimotor network was identified using the ICA-JADE method during the probe stimulus S3. In general, authors of [15] only identified the visual network with a clear temporal localization. Unlike our approach where we used all trial data stacked vertically to emphasize temporal similarities, their analysis is performed by stacking the across-trial averaged data of multiple subjects side by side emphasizing spatial similarities.

The results of the DMD-based approach showed a clearer temporal dynamics than that of the PCA-based approach with comparable spatial networks.

### Methodological considerations

There were many challenges during the design process of this methodology. First, the number of components to consider per window was challenging since it is hard to predict the exact number of brain networks that are active during a specific task. In this work, we applied the DIFFIT technique to identify the number of components needed for the button press and the left-hand movement and fixed the number of components to 2 per interval for the more complex tasks (the simulated and the motor task). Since both PCA and DMD are based on singular value decomposition at their core and since we are following a fixed sliding window approach, many short-term and long-term networks might not be captured. In addition, non-linear evaluations of the brain networks remain challenging and the methods used might not capture all the active networks during tasks. The choice of the window size is also challenging because we do not know in advance how localized in time is a network configuration. Therefore the sampling performed during the computation of the dFC measure needs to be carefully chosen. In addition, when applying the multi-window analysis depicted in figure 3, we need to choose the best number of dFC samples to use noting that we chose it here arbitrarily. Concerning the threshold values, it was challenging to choose the correct value as it was different for the three tasks. This may be due to the strength of the activity of these networks and MEG data which is extracranial collected is not able to capture them all. Nevertheless, we presented statistically significant network values to overcome this difficulty although the statistical analysis across trials, where we used the two-sided t-test with 95% confidence interval with Bonferroni corrections might not be the best approach as well. The null hypothesis statistics were derived from all the windows which may include other non-essential information. Other approaches might be needed to extract more accurate spatial networks such as ANOVA [36]. It is also worth noting that we didn’t use trial averaging except for the case of HCP data (that contained a very large number of subjects with many trials) and argued the need to average them to reduce computational cost and resources needed. For the other datasets, all the trials were used to allow the PCA and the DMD to statistically derive meaningful brain dynamics. One limitation of the method is that when the duration of the brain network activity is short and window-based approaches miss capturing them. In the future, we intend to solve these problems using an incremental version of the methodology using incremental DMD and instantaneous dFCs and constrained dictionary learning approaches such as LANDO [37] to enhance resolution and extract all the dynamics in the data. The methodology described in this paper was applied to cognitive tasks, however, it would be interesting to apply the pipeline to resting-state experiments during which the brain dynamically evolves by shifting from one brain configuration to the other [7]. In addition, the methodology can be tested for epilepsy analysis where the brain often evolves in a specific pattern shifting between different brain configurations since preictal, seizure onset, ictal and postictal intervals showed a specific sequence of states that were repeatable in subjects with epilepsy [38].

## 5. Conclusion

Extracting dominant brain dynamical networks has been extensively studied lately to link tasks to specific brain configurations and network transitions. In this paper, we described a framework for extracting spatial brain networks and their temporal evolution during tasks through multi-trial multi-subject analysis. Experiments performed on synthetic and real MEG data during three motor tasks (simple button press, left hand movement, working memory) showed consistency between the PCA-based and the DMD-based approaches as well as with other studies that employed the same datasets. The main objective of this paper is to design a framework for automatic identification of brain states. In particular, the DMD based approach showed better temporal resolution than the PCA-based approach. In terms of spatial networks identification, both methods had similar performance knowing that both methods make use of the singular value decomposition (SVD) [32] method to learn the discriminative network configurations.

## Acknowledgements

This work was supported by the Institute of Clinical Neuroscience of Rennes (Projects named EEGCog and EEGNET3). The authors of this work declare no competing financial interests and no conflicts of interest.

## Supplementary Material

Supplementary Material is available.

## Data and Code Availability

The HCP data are available on https://www.humanconnectome.org/software/hcp-meg-pipelines. The other datasets are available upon request.

The MATLAB codes are available on https://github.com/Hmayag/Task-Related-DMD-Analysis

