## Supplementary Material for "Analysis of task-related MEG functional brain networks using dynamic mode decomposition"

### I. Supplementary Material

#### 1.1 Functional Connectivity Measures

##### 1.1.1 Amplitude Envelope Correlation

Consider a real valued signal $s(t)$. Complexifying the signal using Hilbert transform $H\{.\}$, $s(t)$ can be written as:

$$\tilde{s}(t) = s(t) + \mathrm{i}\, H\{s(t)\} \quad (1)$$

The analytic signal $\tilde{s}$(t) admits the following form:

$$\tilde{s}(t) = \mathrm{A(t)}.e^{i\varphi(t)} \;\; \mathrm{A(t)} \in \mathrm{R},\; \mathrm{A} >= 0, \varphi(t) \in \mathrm{R} \quad (2)$$

where A(t) is the instantaneous amplitude of $s(t)$ and $\varphi(t)$ is the instantaneous phase, modulo 2π. The instantaneous amplitudes are then used to compute the Pearson correlation coefficient between all pairs of channels resulting in the AEC matrix [29] [30]. The approaches described here use a window-based approach which usually fails to extract meaningful information from small time windows especially from noisy measurements. Therefore, the instantaneous version of the amplitude envelope correlation was developed and tested [9].

##### 1.1.2 Dynamic Phase Locking Value

The phase-locking value (PLV) is used to measure the synchrony between two signals using the phase differences. PLV was formulated to measure inter-trial variability by computing the average phase difference between trials. The PLV measure was extended to quantify the locking at a specific frequency f. A complex wavelet at the predefined frequency is used to filter the considered signals and the Hilbert transform is applied to compute the phase $\varphi_i$(f,t) of each signal. The PLV between N trials for a specific frequency f at time t between two signals is given by:

$$PLV(t,f) = \frac{1}{N}\left|\sum_{n=1}^{N} e^{j\Delta\varphi(t,n)}\right| \quad (3)$$

where $\Delta\phi(t,f,n) = \phi_1(t,f,n) - \phi_2(t,f,n)\ (mod\ 2\pi)$, $\phi_1(t,n)$ and $\phi_2(t,n)$ are the phase of the two signals at time t and trial n. The intuition behind PLV is that when the phase differences remain constant across trails, the signals remain locked as they vary. PLV is close to 1 when the differences remain constant and close to 0 otherwise.

#### 1.2 Source Separation Techniques

##### 1.2.1 Principal Component Analysis

The principal component analysis is a technique that finds m dimensional subspace where the basis vectors correspond to the maximum variance in the vector space of a m dimensional data vector $x = [x_1, x_2\ , \ldots, x_m]$. $x$ is projected into the new subspace to find weights that describe the contribution of each vector. The data is first treated to remove the mean and then normalized. The basis of the PCA can be computed by finding the eigenvectors of the matrix $C_x$ defined by

$$S_x = \sum_{i=1}^{m}(x_i - \mu)(x_i - \mu)^T \tag{4}$$

where μ is the mean of $x$ and $x_i$ is the i[th] data in the vector $x$.

$$C_x = S\Lambda S^{-1} \tag{5}$$

where $\Lambda$ is a diagonal matrix containing the variances of the principal components. PCA learns orthogonal basis directions [31].

Singular value decomposition (SVD) [32] of a $m \times n$ matrix A decomposes the signal into three matrices: a unitary real or complex matrix $U$ $(m \times m)$, rectangular diagonal non-negative real matrix Σ $(m \times n)$, and a unitary real orthonormal matrix $V$ $(n \times n)$. Therefore,

$$A = U\Sigma V^* \tag{6}$$

The diagonals of the matrix Σ are ordered in decreasing order allowing an easy way to reconstruct the original matrix A using the reduced basis. For good approximation, the reduction dimension r to can vary based on the application and the data.

In this study, we employ SVD technique for PCA calculation to extract dominant networks across trials and across patients during simple and complex tasks. The resulting components comprise the spatial network configurations with their respective time dynamics. The extracted dynamic functional connectivity time series data is concatenated vertically for all patients to emphasize temporal similarities and then the resulting spatial components for each temporal component are averaged across the patients to find the dominant spatial networks.

#### 1.2.2 Dynamic Mode Decomposition

Dynamic mode decomposition (DMD) [18] extracts dynamic information purely relying on data. The assumption is based on the existence of a low-dimensional subspace from which the data can be approximately reconstructed linearly. Using experimental snapshots of the data, DMD tries to make sense out of the inherent structure of the data instead of learning models.

A nonlinear continuous dynamical system can be represented using an ordinary differential equation of the form:

$$\frac{dx}{dt} = f\big(x(t)\big) \tag{7}$$

where x(t) ∈ $R^n$ represents the state vector at time t and the function f represents the dynamics function of x. DMD method approximates locally the non-linear dynamical system linearly:

$$\frac{dx}{dt} = Fx \tag{8}$$

whose solution can be written as:\

$$x(t) = \sum_{k=1}^{n} \phi_k e^{\omega_k t} b_k = \Phi e^{\Omega t} b \tag{9}$$

where $\omega_k$ and $\varphi_k$ denote the eigenvalues and the eigenvectors of the matrix $F$, and $b_k$ is the projection of the initial condition $x(0)$ on the eigenvector basis. In the discrete domain, equation (8) can be sampled every Δt and can be written as:

$$x_{k+1} = Ax_k, k = 0,1,2,\dots,m \tag{10}$$

where $A = e^{F\Delta t}$ and m is the number of snapshots collected. The discrete solution of equation 10 is:

$$x(t) = \sum_{j=1}^{n} \phi_j \lambda_j^k b_j = \Phi \Lambda^k b \quad (11)$$

where $\lambda_j$ and $\varphi_j$ are the eigenvalues and the eigenvectors of the matrix $A$ and $b$ is such that $x_0 = \Phi b$. Therefore, the DMD algorithm finds in the least square sense, a low-rank eigendecomposition of the matrix A by minimizing:

$$||x_{k+1} - Ax_k||_2 \quad (12)$$

for $k = 0,1,\ldots,m-1$.

Let $x_k \in R^n$ denote the measurements from n channels at instant $k$. The matrix $X$ resulting from horizontally stacking these measurements has a dimension $nxm$ as shown below.

$$X = [x_0, x_1, \ldots, x_{m-1}] \quad (13)$$

Construct $X'$ which contains one time shifted version of the data as :

$$X' = [x_1, x_2, \ldots, x_m] \quad (14)$$

Therefore,

$$X' = AX \quad (15)$$

The solution of the least squares problem is:

$$A = X'X^{\dagger} \quad (16)$$

where † is for the Moore-Penrose pseudoinverse. The eigenvectors and the eigenvalues of the $A$ correspond to the modes and eigenvalues of the DMD. Note that, if the size of the vector n is large, the matrix $A$ is hard to analyze and the pseudo-inverse becomes computationally expensive. The DMD algorithm instead computes a lower dimensional representation of the data in the matrix $X$ by finding an approximation $\tilde{A}$ of $A$. The algorithm is shown below where * denotes the complex conjugate of the matrix, the left singular vectors of $X$ are $U_r \in R^{nxr}$, the right singular vectors of $X$ are $V_r \in R^{mxr}$,the singular values of $X$, $\Sigma_r \in R^{rxr}$ , $r$ is the number of singular values kept where $r < min(m,n)$, the eigenvector of $\tilde{A}$ are $W \in C^{rxr}$, the eigenvalues of $\tilde{A}$ are $\Lambda \in C^{rxr}$, the DMD modes $\Phi \in C^{nxr}$ relating channels to the modes. The result is a DMD tuple $(\lambda, \phi, \varphi)$ defined by the eigenvalues $\lambda_i$, the eigenfunctions $\phi = e^{\Omega t}$, and the modes $\varphi$. Oscillatory modes are represented by complex conjugate eigenvalue pairs. Therefore, the model captures at most $r/2$ distinct frequency components.

**DMD Algorithm**

Result: A, DMD modes, DMD eigenfunctions, DMD eigenvalues

Input: A

1. Find the singular value decomposition (SVD) of the matrix $X = U\Sigma V^*$ and reduce the rank (by taking top r eigenvalues and eigenfunctions) of the matrices to approximate it to $\widetilde{X} = U_r \Sigma_r V_r^*$.
2. Compute $A = X'V_r\Sigma_r^{-1}U_r^*$. Define $\widetilde{A} = U_r^* A U_r$ . Therefore, we have $\widetilde{A} = U_r^* X' V_r \Sigma_r^{-1}$.
3. Eigen decompose the matrix $\widetilde{A}$ using $\widetilde{A}W = W\Lambda$ where W contains the eigenvectors and $\Lambda$ is the diagonal matrix containing the DMD eigenvalues $\lambda_i$.
4. Compute the DMD modes $\Phi \approx X'V\Sigma^{-1}W$. Each column $\phi_i$ of $\Phi$ is a DMD mode corresponding to each eigenvalue $\lambda_i$.

5. Reconstruct the signal as a composition of coupled spatio-temporal modes as: $\hat{X} = \Phi e^{\Omega t} b$ where $\Omega = \frac{\log(\Lambda)}{\Delta t}$, $\Delta t$ is the sampling duration, and b is computed from initial conditions such that $x_0 = \Phi b$.

---

#### 1.2.3 Time Delay Embeddings

DMD was originally developed for the case where the state dimension n is much greater than the number of observations or snapshots m. However, if $m >> n$, then DMD cannot capture the nonlinear dynamics fully (cite 28/29 from Koopman Operator Applications in Signalized Traffic Systems). Hankel Time delay embedding is a method to overcome this deficiency where states are augmented by $h$ measurements with past history. The reconstruction of the states depends on the lag time between snapshots $\delta$ and the embedding dimension $h$. Here, $\delta = k\Delta t$ where $k$ is any integer and generally $k = 1$. The parameter h is chosen such that $hn >> (m - h)$. The Hankel matrix is formed

$$\tilde{X} = \begin{pmatrix} x_1 & x_2 & \cdots & x_m \\ x_2 & x_3 & \cdots & x_{m+1} \\ \vdots & \vdots & \ddots & \vdots \\ x_h & x_{h+1} & \cdots & x_{h+m-1} \end{pmatrix} \quad \tilde{X}' = \begin{pmatrix} x_2 & x_3 & \cdots & x_{m+1} \\ x_3 & x_4 & \cdots & x_{m+2} \\ \vdots & \vdots & \ddots & \vdots \\ x_{h+1} & x_{h+2} & \cdots & x_{h+m} \end{pmatrix}$$

DMD is applied on the Hankel matrix $\tilde{X}$ and $\tilde{X}'$ (instead of X and X′) to approximate the Koopman tuple (λ,ϕ,φ) The columns of V represent the principal components of the trajectory. Next, r components are chosen to approximate the system.

In this study, DMD with h time delay embedding is used to further smooth the temporal components and find underlying dynamic modes from the data. For each smoothed temporal component, the complex DMD modes are used to analyze the spatial networks where the real, imaginary, and magnitudes are considered. Note that, DMD modes come in pairs since conjugate complex pairs are required to reconstruct the real signals with the complex components.

### II. Supplementary Results

#### **2.1.1 PCA vs DMD on toy data**

In this section, PCA and DMD are applied on simple synthetic oscillatory data to demonstrate how DMD is superior to PCA in terms of phase variability between measurements specifically for temporal latency jitters introduced naturally due to the variability of trials and subject responses.

Consider 1.6-second long 8-channel synthetic time series, denoted by $Y_i$ where i = 1,2,...,8 sampled at 250 Hz in which the first three channels oscillate at 6.6Hz and exhibit some random phase lag between each other, and where channels 4,5,6,7,8 oscillate at 12.5 Hz in phase as shown in figures S1A and S2A. The amplitudes of the signals are 0.5 and the data is contaminated with random noise of magnitude 0.05 (SNR = 20).

Applying PCA on Y with 3 principal components extracts the temporal components that are illustrated in figure S1B, along with their corresponding spatial components in figure S1C. The first principal component, PC1, captures the 12.5Hz oscillations whose spatial principal component is high in channels 4-8 as seen in S1C in red. On the other hand, the second component, PC2, captures the 6.6 Hz oscillations of channels 1, 2, and 3 whose spatial principal component vector shows that even channels 5-8 rely on PC2 to reconstruct the initial data. However, these oscillatory components are not sinusoidal in nature although the data originally is sinusoidal. The reason for this behavior is the phase lags between the three channels. Finally, PC3 illustrated as the yellow time series in figure S1B, also plays a role in reconstructing the original signal. The mean square error of reconstructing the signals Y using PCA with three components was 0.0613.

On the other hand, DMD is applied on the signals Y using 6 modes (since DMD modes come in pairs that are homologous to the 3 components used in the PCA) and 20 time-delay embeddings (chosen arbitrarily). Figure S2B shows the real part of the time dynamics of modes 1,3, and 5, and their corresponding imaginary parts are shown in figure S2C. Note that, the real parts of the time dynamics of modes 2,4, and 6 overlap the real parts of modes 1,3,and 5. The corresponding spatial components that were extracted by using the magnitude of the DMD mode matrix $\Phi$ are illustrated in figure S2D where mode 1 is active only in channels 4-8, while mode 2 explains the activity in channels 1-3. Finally, the spatial components of mode 3 are randomly expressed in all the channels knowing that their temporal dynamics are close to zero (as seen in figure S2B yellow signal), they have no significance. DMD overcomes the problem of the phase lags between signals and clearly distinguishes different oscillatory components by expressing their dynamics in different modes.

Finally, figure S3 shows the phase analysis of the DMD mode matrix $\Phi$. As discussed above, mode 1 summarizes the dynamics of channels 4-8, while mode 2 explains channels 1-3.

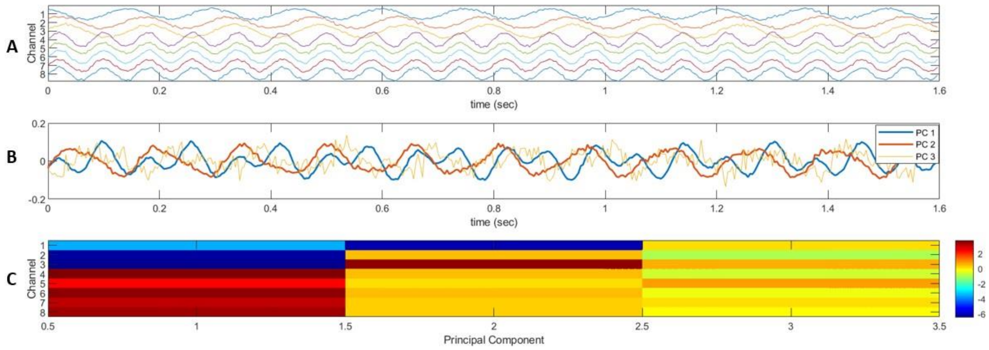


Figure S1: PCA spatial and temporal components applied on synthetic data

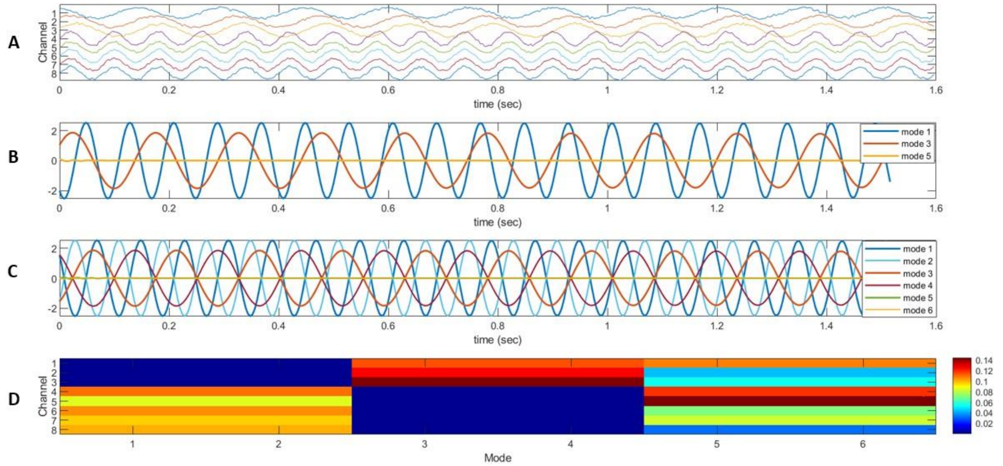


Figure S2: DMD spatial and temporal components applied on synthetic data

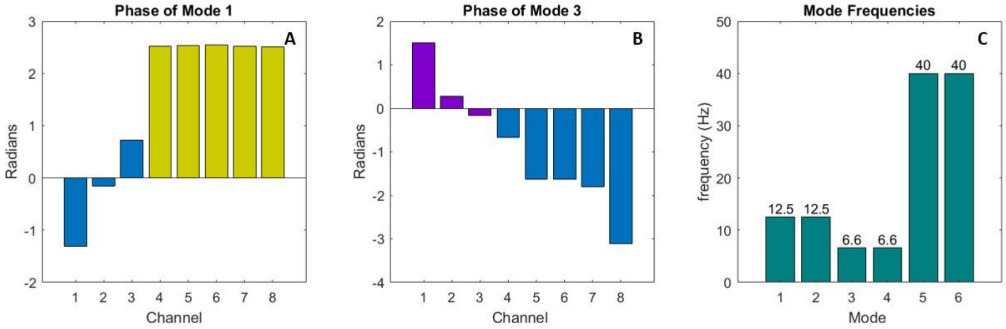

Figure S3: DMD phase analysis of the synthetic data

While channels 4-8 oscillate in phase with each other in mode 1 ( yellow bars in figure S3A), channels 1-3 oscillate at different phase lags in mode 2 (purple bars in figure S3B). However, the magnitudes of the DMD modes shown in figure S2D clearly separates the activity in different channels even despite the phase lags. Therefore, the phase information is preserved as well and can be evaluated if needed. Finally, the mode frequencies are extracted from the eigenvalues of the DMD operator that capture the exact frequency values of the original data: 12.5 Hz for mode 1 and 6.6 Hz for mode 3. Modes 5 and 6 were found to oscillate at 40 Hz that model the noisy components roughly and which is ignored in the analysis as seen in figure S3C. The mean square error of reconstructing the signals Y using DMD with six modes was 0.0107 which is less than that of the PCA.

#### 2.1.2 Simulated EEG data

This dataset is composed of simulated human EEG time series based on an interconnected Neural Mass Model (NMM) that generates coherent oscillations in different brain structures similar to real EEG during a picture-naming task [22]. The neural mass model constructs 66 regions of interest (ROI) from the standard anatomical parcellation of the Desikan-Killiany atlas. The data correspond to a 2000msec long time-series where a picture-naming task was simulated by taking a baseline activity and then inputting six simulated networks consecutive in time from 1000 to 1535 msec that were activated to mimic the picture naming task by tuning the parameters of the NMM to generate the desired background and gamma activity as shown in figure S4. Using a sampling rate of 1024 Hz, the gamma band activity [30-40 Hz] was used as it is the most relevant frequency band related to the cognitive tasks. 20 subjects with 100 trials each were simulated by varying the connectivity and the input noise uniformly resulting in 2000 2-second long simulations. A detailed description of these simulations can be found in [23].

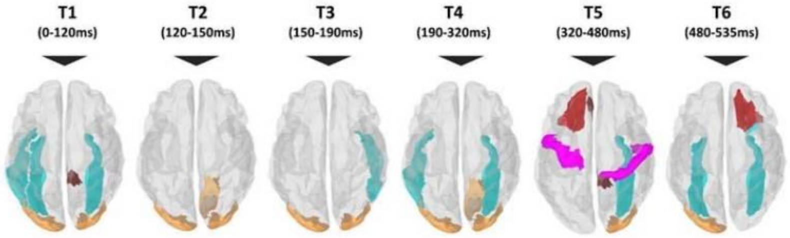


Figure S4: Simulated network configuration (adapted from [23]).

#### 2.1.3 Results of the workflow on Simulated EEG data

For the synthetic data, since the data is long and the dynamics shifts rapidly, we employed 2 principal components for each interval which is valid since evaluations are done per interval and by design, each interval has a single network configuration. The workflows depicted in figures 1, 2, and 3A are applied on this synthetic data.

The networks were generated by varying parameters of the neural masses as desired in each interval $T_i$. As mentioned earlier, the dataset generated is source time series therefore the beamforming step is skipped and the dynamic functional connectivity is computed directly. The instantaneous phase-locking values are computed FOR each subject across trials according to equation (3) resulting in a 66x66x2049 dFC matrix. First, the inter-subject analysis at the six intervals $T_1$ to $T_6$ is performed. The dFC time series data of all the subjects are concatenated vertically and workflow 1 (shown in figures 1 and 2) is applied to extract dominant spatial components using PCA and the DMD techniques. The t-test with Bonferroni correction is applied to find only the significant values with 95% confidence.

Two components are used for the PCA while r is set to 4 in the case of the DMD. Note that, with $r = 4$, DMD extracts two networks since DMD modes come in pairs as discussed earlier. We employ the magnitudes of the complex modes. Figure S5 shows the 2 dominant networks using the PCA method for the six intervals applied separately across all subjects. Similarly, figure S6 shows the results for the DMD method. Both the PCA and the DMD approaches to capture the original chosen setup in all the intervals.

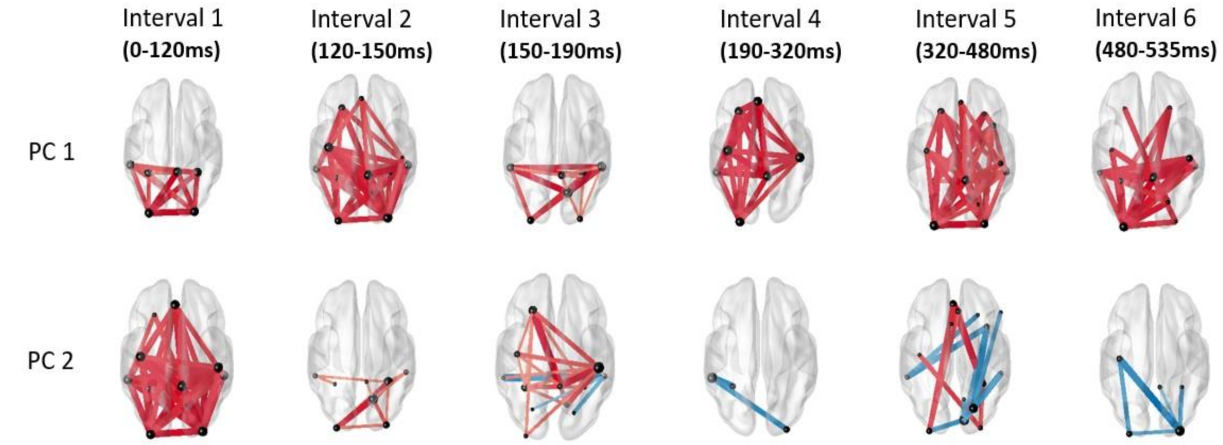


Figure S5: Principal components in the six intervals of the simulated dataset using PCA

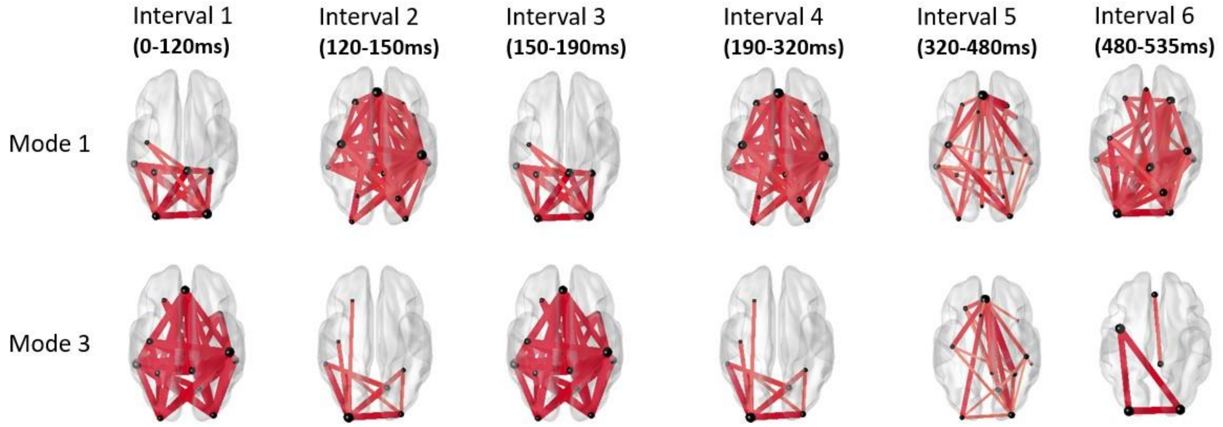


Figure S6: Principal spatial modes in the six intervals of the simulated dataset using DMD

Next, we apply the sliding window approach across the 2-second interval using a window size of 50 samples ( 49 milliseconds) with 49 sample overlap (step size is 1 sample) to extract for each window the two dominant networks for both PCA-based and DMD-based approaches. The resulting instantaneous components are collected in a single matrix.

Finally, PCA with 9 components (determined using DIFFIT) is applied to extract the dominant networks and the corresponding time dynamics across the whole experiment. The results for the PCA-based approach are shown in figure S7. It can be seen that PC1 which exhibits activity in the temporal, occipital, and prefrontal, is active in two major intervals: 1000-1100 msec,1300-1400 msec, and around 1500 msec homologous to the 1$^{st}$, 4$^{th}$, and 5$^{th}$ interval. On the other hand, PC3 shows activity only in the interval 1450-1550msec which is homologous to the 6$^{th}$ interval.

The results for the DMD-based approach are shown in figure S8 where many of the networks are identified and their corresponding time evolutions are comparable with the original network configurations shown in figure S4.

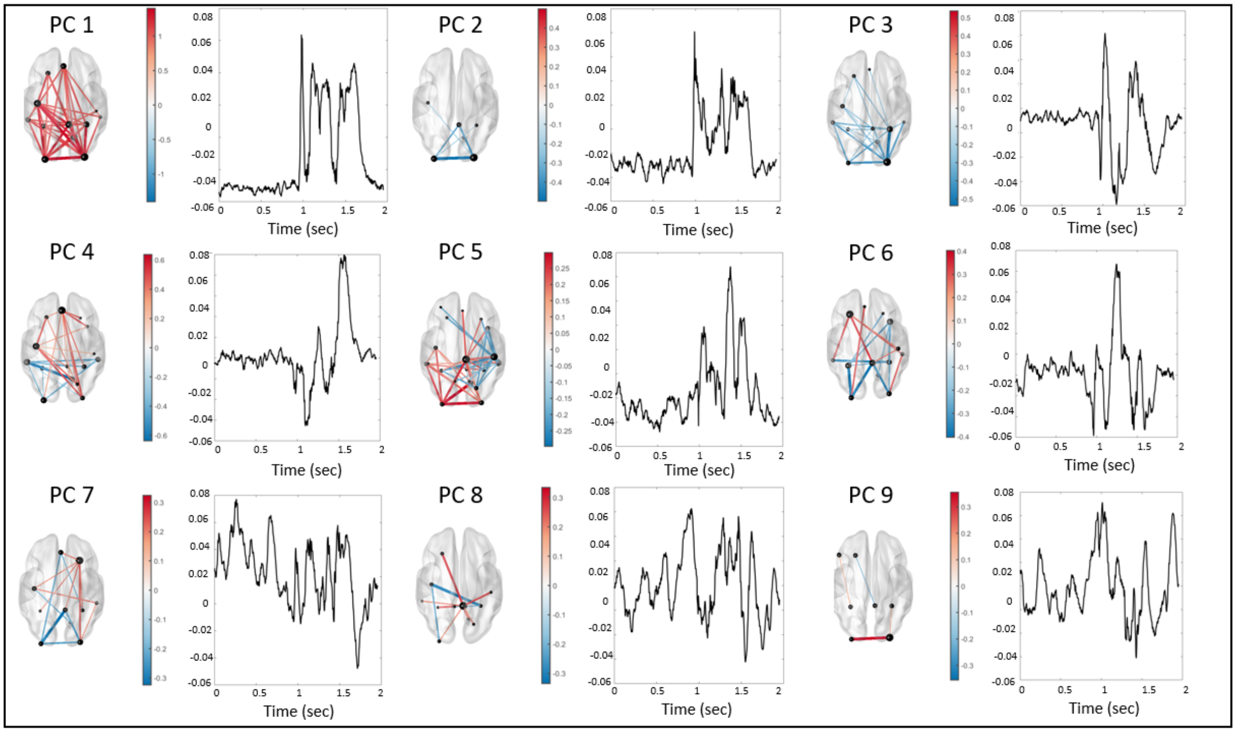


Figure S7: The principal spatial components across all intervals using the PCA-based networks

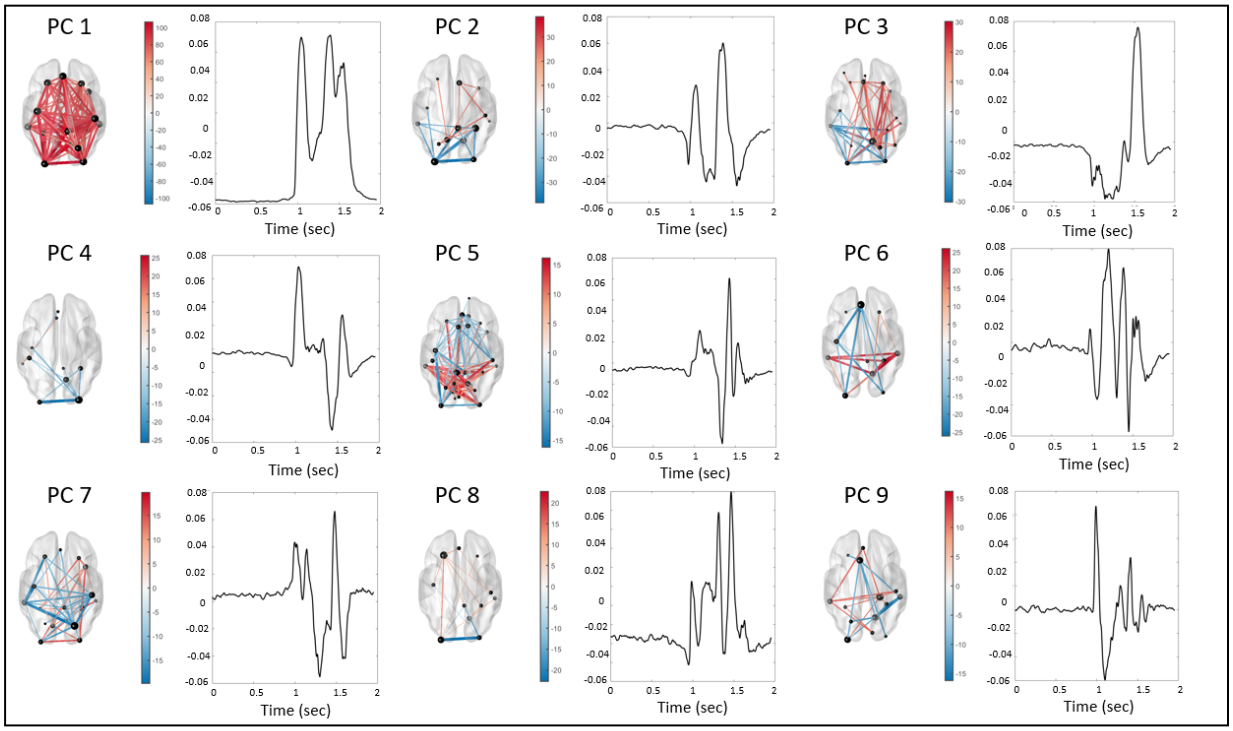


Figure S8: The principal spatial components across all intervals using the DMD-based networks

### III. Discussion

In this study, our aim was to learn dominant spatial configurations during specific tasks and identify the temporal timestamps of each. The main aim was to introduce the DMD approach and compare it to the PCA methodology. Since there exists inter-trial variability during the experimental acquisition of the data, we first compared DMD and PCA on a simple toy example where phase distortions were introduced between different time series and we compared their dominant spatial and temporal components. DMD with time delay embeddings was superior to the PCA method in various ways. First, DMD can compress the dynamics in multi-channel data spatially, temporally, and spectrally, especially for oscillatory data. Second, DMD compresses phase lagged dynamics of similar spectral components into a single mode while keeping track of the phase information of the channels. The advantages of the DMD can help overcome the limitations of the PCA specifically in multi-trial and multi-subject analysis, in which temporal variability exists. In addition, since multiple trials are generally averaged to increase the SNR value, DMD provides the necessary mechanism to overcome this problem.

Next, we evaluated the proposed methodology on synthetic data to validate our approach before testing it on real MEG data. Here the data was simulated with known ground truth and the results of our methodology using the PCA-based and the DMD-based approaches were compared. Both the PCA and the DMD approaches provided similar spatial configurations, however, the DMD approach’s temporal components were smoother. One limitation here was the lack of quantitative evaluation method since the study was performed on the data that was already simulated with the shown simulated networks shown in figure S4.
